# Progranulin loss induces mitochondrial dysfunction and ferroptosis in human cerebral organoids

**DOI:** 10.64898/2025.11.29.691344

**Authors:** Carolina M. Camargo, Maria Camila Almeida, Bárbara Mejía-Cupajita, Inyoung Hwang, Mohamed A. Faynus, Daniel C. Carrettiero, Kenneth S. Kosik

**Affiliations:** Neuroscience Research Institute, Department of Molecular, Cellular and Developmental Biology, University of California, Santa Barbara, Santa Barbara, CA 93106, USA; Center for Stem Cell Biology and Engineering, Neuroscience Research Institute, University of California, Santa Barbara, Santa Barbara, California, CA 93106, USA

## Abstract

Loss-of-function mutations in the granulin (*GRN*) gene cause frontotemporal dementia when the mutations are heterozygous and neuronal ceroid lipofuscinosis, a lysosomal storage disease, when homozygous. While it is well established that disease-causing *GRN* mutations decrease progranulin (PGRN) levels, leading to neurodegeneration, the cellular and molecular mechanisms underlying these conditions remain poorly understood. In this study, we utilized human induced pluripotent stem cell (iPSC) derived forebrain organoids to investigate the impact of PGRN homozygous deficiency on neuronal and glial cell populations. Through single-cell RNA sequencing, we identified robust downregulation of the mitochondrial oxidative phosphorylation pathway in PGRN KO organoids. In line with these results, PGRN KO organoids showed decreased mitochondrial respiration. Furthermore, our study demonstrated that PGRN loss induced increased levels of reactive oxygen species (ROS), lipid peroxidation and iron accumulation. Finally, we observed increased vulnerability to ferroptotic cell death in PGRN KO organoids. Our findings suggest that mitochondrial dysfunction and impaired responses to oxidative stress are early manifestations of PGRN loss, and offer insights into the molecular mechanisms driving neurodegeneration caused by PGRN deficiency.

Graphical Abstract
Proposed molecular mechanisms that lead to ferroptosis of PGRN KO cells. Created with BioRender.com.

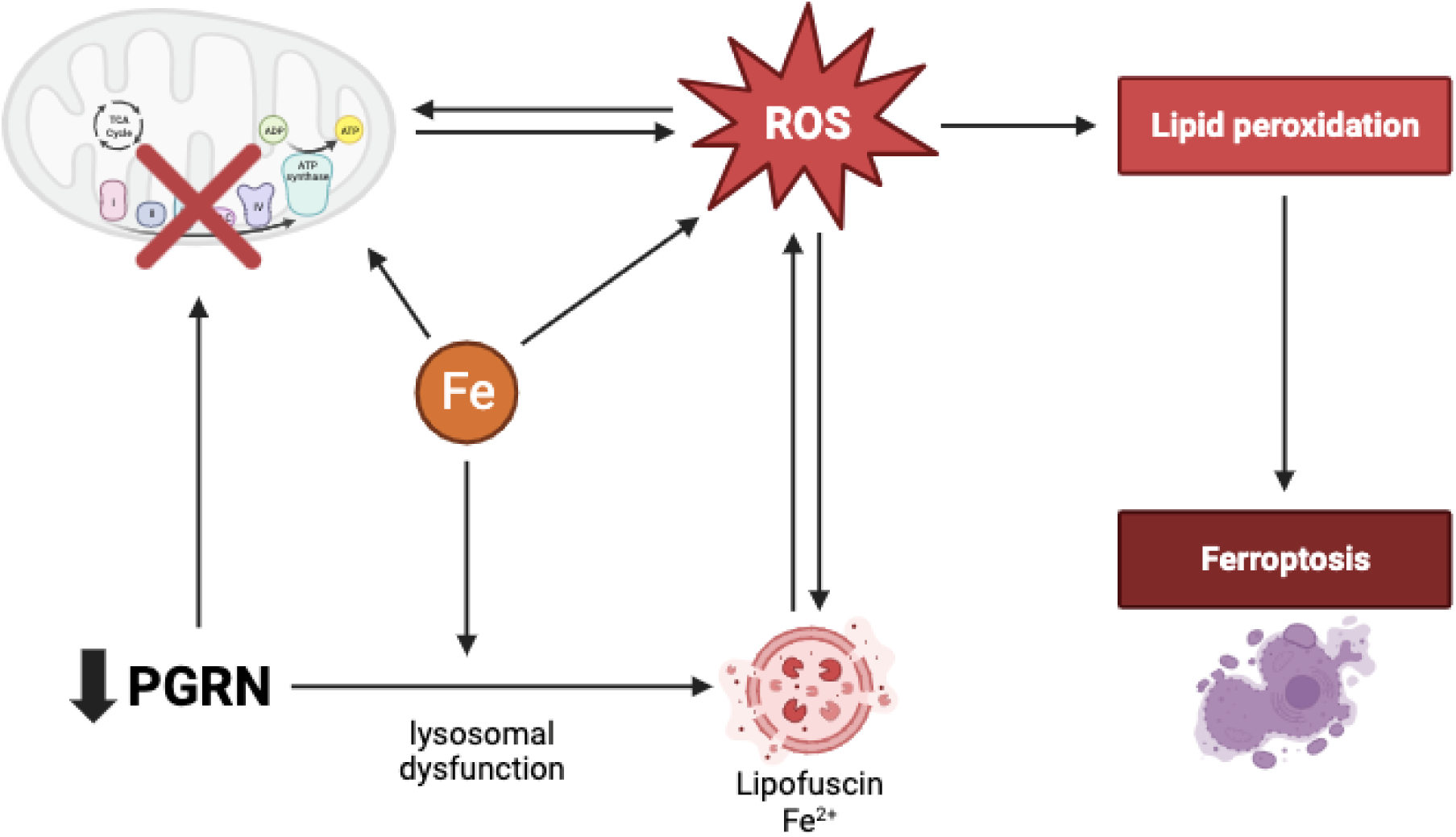

## Introduction

Neurodegenerative diseases are characterized by progressive neuronal loss and chronic inflammation, processes that are strongly influenced by dysregulated lysosomal function and impaired neurotrophic signaling. Among the key regulators of these pathways is progranulin (PGRN). Progranulin is a multifunctional, secreted and intracellular protein expressed in neurons, microglia, endothelial cells and astrocytes. In the central nervous system (CNS), PGRN acts as a neurotrophic factor, regulates lysosomal homeostasis and modulates neuroinflammation (1–3).

Mutations in *GRN*, which encodes PGRN, cause neurodegeneration in a gene-dose–dependent manner - heterozygous mutations lead to frontotemporal dementia (FTD) with TDP-43 inclusions (4–6), whereas homozygous loss causes CLN11, a rare form of neuronal ceroid lipofuscinosis (NCL) (7,8). Most *GRN* mutations are loss-of-function variants that reduce or abolish PGRN expression through nonsense-mediated mRNA decay (6), and, accordingly, progranulin AAV gene therapy treatments have provided early evidence of therapeutic potential (9,10). Although the clinical manifestations of these diseases are distinct, they share some common pathological features such as lipofuscin accumulation, impaired lysosomal function, neuroinflammation and gliosis (11–13). Additionally, mutations in *GRN* are associated with enhanced susceptibility to Parkinson’s disease (PD) (14), amyotrophic lateral sclerosis (ALS) caused by repeat expansions in the *C9ORF72* (15) and Alzheimer’s disease (AD) (2,16), and reduced levels of PGRN are associated with accumulation of other proteins besides TDP-43, such as TAU and α-synuclein (17,18). In spite of the clear and important role of PGRN in neurodegeneration, a comprehensive picture of its cellular and molecular functions remains incomplete.

Human induced pluripotent stem cell (iPSC)–derived brain organoids have emerged as a powerful tool for modeling neurodegenerative diseases, capturing cell-type diversity and three-dimensional architecture that more closely reflect the human brain than traditional two-dimensional cultures, and enabling the examination of early pathological events preceding overt neurodegeneration.(19,20). PGRN deficient human iPSC-derived neurons and 3D co-cultures of neurons and astrocytes recapitulate some pathological hallmarks of FTD-GRN and NCL, such as mislocalization and hyperphosphorylation of TDP-43, enlarged lysosomes and lysosomal lipofuscin accumulation (21–24). However, many 2D models do not display aspects of TDP-43 pathology seen in FTD-GRN patients, which suggests that PGRN induces pathology via non-cell autonomous mechanisms (21,25,26). Indeed, *GRN* deletion in only microglia or neurons is not sufficient to cause CLN11-like pathology (27,28), and PGRN-deficient astrocytes and microglia can induce toxicity in control neurons (26,29,30). These findings indicate that the combined contributions of multiple cell types are likely required for the full manifestation of PGRN deficiency phenotypes.

In this study, we used forebrain organoids derived from *GRN* knockout and control iPSCs to interrogate the cellular consequences of progranulin deficiency. Through single-cell RNA sequencing (scRNA-seq) we revealed significant downregulation of genes related to mitochondrial oxidative phosphorylation in PGRN KO organoids compared to controls. Consistent with these findings, PGRN KO organoids exhibited reduced mitochondrial respiration, increased susceptibility to ferroptotic cell death, and hallmark features of ferroptosis, including reduced GPX4 expression, iron accumulation, and increased lipid peroxidation. Together, our findings uncover a previously unrecognized link PGRN homozygous deletion in association with both mitochondrial dysfunction, and ferroptotic cell death.

## Results

### Generation and single-cell RNA-seq of PGRN KO and isogenic control cerebral organoids

Organoids were generated for two human iPSC lines – PGRN KO (*GRN^-/-^*), and an isogenic control – Ctrl (*GRN^+/+^*). Both iPSC lines were generated in the WTC11 iPSC background, which was mutagenized to generate the PGRN KO iPSC (21) (Supplementary Figure S1A). Human dorsal forebrain organoids (hCO) and ventral forebrain organoids (hVO) were differentiated following a directed-differentiation protocol (20,31), then fused at 2 months for subsequent scRNA-seq analyses. It has been shown that 6-month organoids present a higher variety of cell types and a larger proportion of neurons than younger organoids (20,31); therefore we chose this timepoint for our studies. Knockout of *GRN* was confirmed on mRNA and protein expression levels (Supplementary Figure S1B, C). Single-cell RNA-seq libraries were generated using the 10X Genomics platform (Figure 1A). Three 6-month old fused organoids from each genotype were sequenced, and following quality control analyses, we obtained a total of 8,975 cells (1496 ± 354 cells per sample).

**Figure 1.**
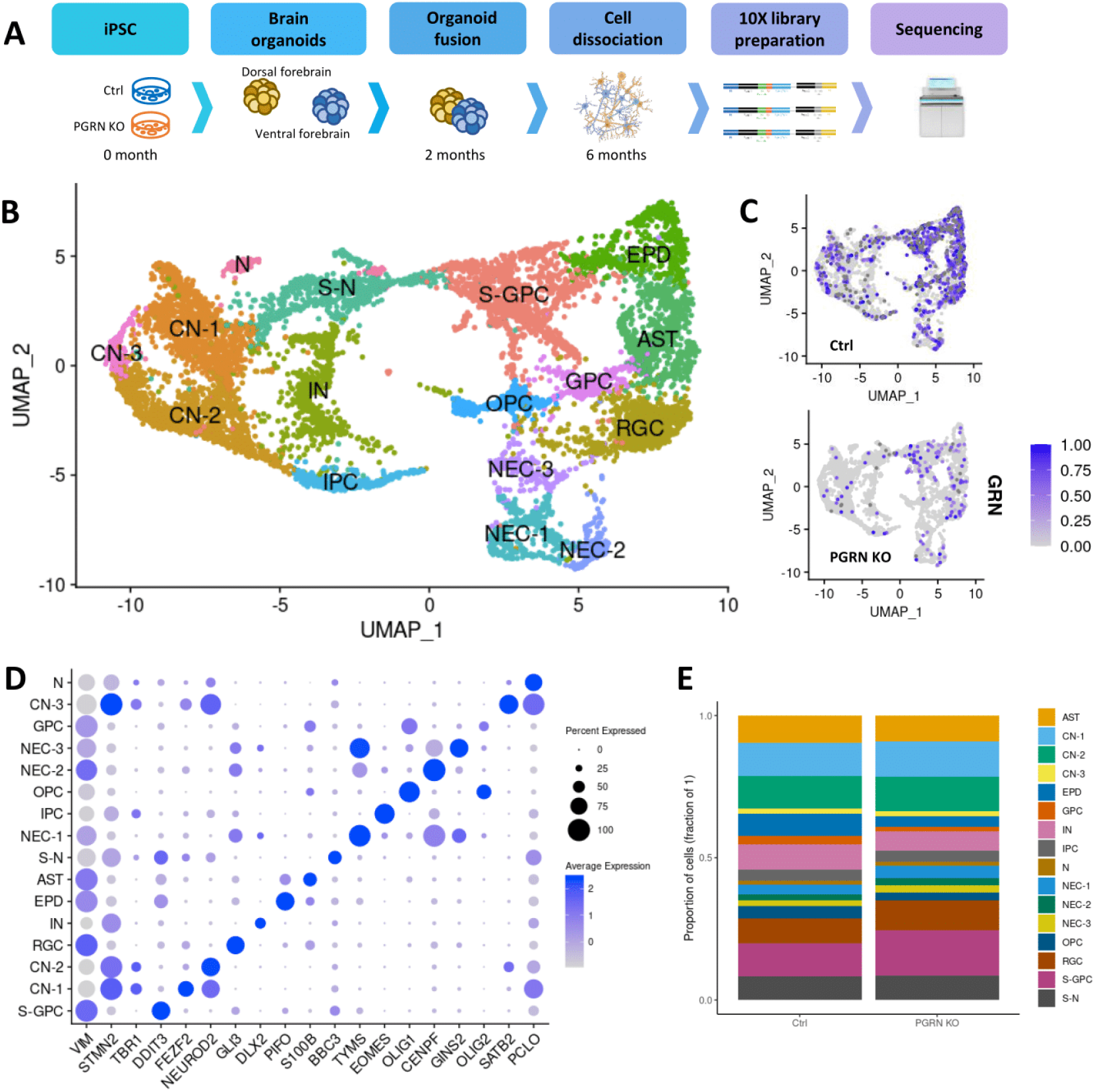
Cellular composition of PGRN KO and Ctrl organoids upon scRNA-seq analysis. A – Schematic illustration of the sample material and scRNA-seq workflow. B – UMAP of 8975 cells from six PGRN KO and Ctrl brain organoids. C – UMAP of PGRN KO and control organoids, with *GRN* expression highlighted in blue. D – Dot plot showing expression of selected markers genes in each cell cluster. E – Stacked bar plot of proportions of major cell types in PGRN KO and control organoids. Statistical significance was calculated by 2-way ANOVA with Sidak multiple comparisons test. CN – cortical glutamatergic neurons, IN – gabaergic interneurons, IPC – intermediate progenitor cells, N – unidentified neurons, S-N – stressed neurons, GPC – glial progenitor cells, S-GPC – stressed glial progenitor cells, EPD – ependymal cells, AST – astrocytes, OPC – oligodendrocyte precursor cells, RGC – radial glial cells, NEC – neuroepithelial cells.

Unsupervised clustering of cells based on their transcriptomic profiles resulted in the identification of 16 major cell types based on expression of unique markers (Supplementary Table S1; Figure 1B, D), which aligns with findings from other scRNA-seq studies on human brain organoids (19,32,33). We identified three clusters of glutamatergic cortical neurons (CN), all expressing the markers *NEUROD6*, *NEUROD2*, *STMN2*, *BCL11B*, *TBR1*, *MAPT* and *DCX*. CN-1 expressed high levels of genes such as *FEZF2* and *SYT4*; CN-2 showed high expression of genes such as *SOX11* (early neuronal marker) and *SATB2*; and CN-3 was characterized by high expression of *SATB2*, among other genes. GABAergic interneurons (IN) expressed high levels of the markers *DLX1*, *DLX2*, *GAD1* and *GAD2*. We also found a population of unidentified neurons (N) expressing the neuronal markers genes *GRIN2B*, *PCLO*, *ANK3* and *MAPT*. Intermediate progenitor neurons (IPC) showed high expression of the markers *EOMES*, *ELAVL4*, *NHLH1* and *PPP1R17*. Neuroepithelial cells (NEC) formed 3 clusters (NEC-1, NEC-2 and NEC-3) and expressed high levels of the cell-cycle related genes *HMGB2*, *TOP2A*, *HELLS* and *CENPF*. Astrocytes (AST) expressed high levels of the canonical markers *NDRG2*, *S100B*, *GJA1* and *AQP4*. Radial glial cells (RGC) formed a cluster characterized by high expression of genes such as *PAX6*, *HOPX*, *TNC*, *VIM*, *GLI3*, *PTPRZ1* and *SLC1A3*. Oligodendrocyte progenitor cells (OPC) expressed genes such as *OLIG1*, *OLIG2*, *DCX1*, *DCX2*, and *PDGFRA*. Glial progenitor cells (GPC) were characterized by intermediate expression of OPC and RGC/astrocyte genes. Stressed glial progenitor cells (S-GPC) expressed intermediate levels of the RGC/astrocyte markers *VIM*, *SLC1A3*, *TNC* and *PAX6*, no expression of OPC genes, and showed high expression of unfolded protein response (UPR) genes (e.g. *DDIT3*, *DDIT4*, *CRYAB*, *P4HB*, *SLC3A2*). A population of cells clustered that expressed intermediate levels of neuronal markers such as *BCL11B*, *STMN2*, *SNAP25* and *SYT4*, as well as high expression of the same S-GPC UPR genes and high expression of other UPR genes such as *HERPUD1* and *BBC3*. We called this population “stressed neurons” (S-N). The unfolded protein response is a stress response pathway that regulates endoplasmic reticulum homeostasis. Similar populations of stressed cells that show upregulation of the UPR have been found in previous single-cell RNAseq organoid studies (19,32), and they most likely represent cells that have been affected by the lack of oxygen supply due to the absence of vascularization in brain organoids (19,32). Finally, we identified a population of ependymal cells (EPD), characterized by high expression of the markers *FOXJ1*, *PIFO* and *DYNLRB2*. *GRN* was expressed in all cell types in our dataset (Figure 1C). All known major cell types were detected in both PGRN KO and Ctrl organoids, with consistent similar numbers across samples (Figure 1E).

### Dysregulation of oxidative phosphorylation system in PGRN KO organoids

To identify dysregulated genes related to PGRN loss, we performed differential gene expression analysis between PGRN KO and Ctrl organoids for each identified cell type (absolute log2FC > 0.25, adjusted p-value < 0.05) (Supplementary Table S2), and then conducted gene set enrichment analysis to identify biological processes and pathways that were overrepresented among the differentially expressed genes (DEG).

We found that the most significant dysregulated pathway of downregulated genes in all cell types was aerobic respiration and respiratory electron transport, which includes genes involved in mitochondrial oxidative phosphorylation (OXPHOS) (Figure 2A; Supplementary Figure S2, Supplementary Table S3). The OXPHOS system is the final biochemical process in the production of ATP, and is composed of the electron transport chain (ETC), a set of five multiprotein enzyme complexes localized in the inner mitochondrial membrane that are encoded by genes in the mitochondrial or nuclear DNA (34). The ETC creates a proton gradient utilized to generate ATP production, and produces electron carriers (e.g. NAD+ and FAD) that are essential for glycolysis and the tricarboxylic acid (TCA) cycle. In our study, all dysregulated genes in this pathway were nuclear-encoded genes that encode components of the complexes I (NADH dehydrogenase), III (cytochrome bc1 complex), IV (cytochrome c oxidase) and V (ATP synthase) of the mitochondrial ETC, and most of them overlapped in all the cell types (Figure 2B, OXPHOS genes in astrocytes and neurons). After finding this major dysregulation of mitochondrial respiration genes in PGRN KO organoids, we then focused the differential gene expression analysis in all neurons (CN-1, CN-2, CN-3 and IN clusters combined) and astrocytes (Figure 2C, D), considering these cell types are heavily affected by progranulin deficiency (2,21,29,30).

**Figure 2.**
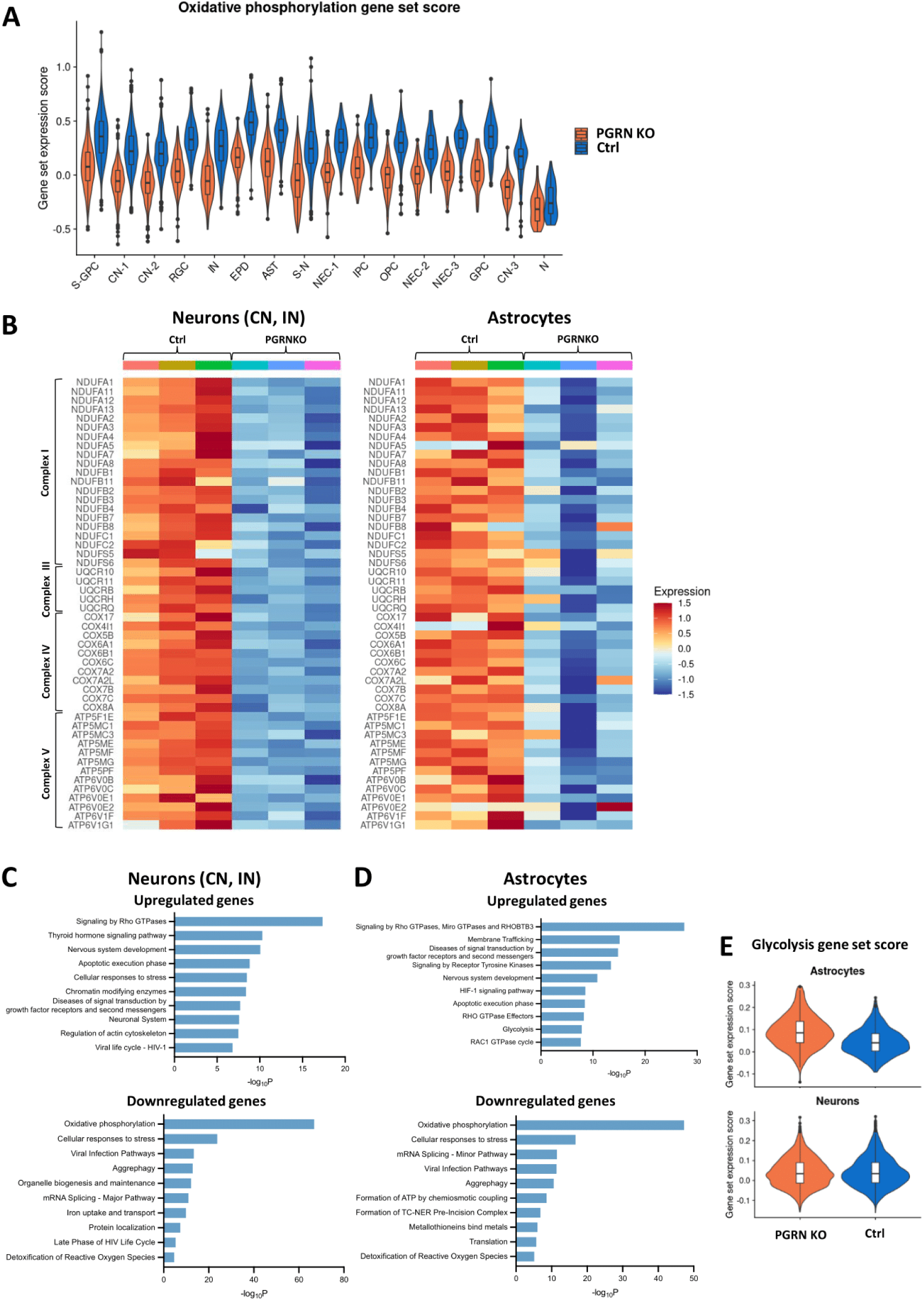
Pathway enrichment analysis and downregulation of OXPHOS in PGRN deficient organoids. A – Gene expression scores for the OXPHOS gene set for all cell types. B – Heatmap of expression levels of DEG involved in OXPHOS in astrocytes and neurons (CN-1, CN-2, CN-3 and IN). C – Bar graph of the top 10 enriched pathways across DEGs between PGRN KO and Ctrl neurons (CN-1, CN-2, CN-3 and IN). DEGs comprised 539 upregulated and 280 downregulated genes. D – Bar graph of the top 10 enriched pathways across DEGs between PGRN KO and Ctrl astrocytes. DEGs comprised 584 upregulated and 225 downregulated genes. E – Gene expression scores for the glycolysis gene set for astrocytes and neurons.

Among the enriched pathways of downregulated genes in both astrocytes and neurons, aggrephagy (Figure 2C, D) stands out. Aggrephagy is the removal of protein aggregates by macroautophagy, and several studies have shown that PGRN deficiency leads to impairment of autophagic flux, which can result in accumulation of TDP-43 (35,36). Mitophagy, the process by which damaged mitochondria are selectively degraded via autophagy, was suggested to be affected in our dataset, as we observed downregulation of several associated genes - including *GABARAPL2, GABARAP, FIS1, JUN, UBB, TOMM7,* and *UBA52 -* in astrocytes and neurons. These findings also align with previous research demonstrating that PGRN deficiency impairs mitophagy (36–40).

Pathway enrichment analysis of downregulated genes also showed that the terms “cellular response to stress” and “detoxification of reactive oxygen species” were enriched in PGRN KO neurons and astrocytes (Figure 2C, D). Among those genes, several involved in the ubiquitin-proteasome system (*UBB, UBA52, PSMA2, ELOB, RBX1, SEM1*) suggested proteasome dysfunction. These pathways also included antioxidant genes that were downregulated in astrocytes (*PRDX5, PRDX1, JUN, PARK7, MT3, TXN, ATOX1, HSBP1, GSTP1*) and neurons (*SOD1, ATOX1, PRDX5, PRDX2, PARK7, MT3, MIF, TXN, HSBP1, MT1F, MT2A*).

The HIF-1 signaling pathway was one of the top pathways of upregulated genes enriched in astrocytes (Figure 2D). These genes included *HIF1A, EGLN1, HK1, VEGFA, PDK1, CDKN1B* and *AKT3*, which are also upregulated in neurons. HIF-1α mediates responses to oxidative stress and hypoxia by translocation to the nucleus and mitochondria, and downregulates OXPHOS to decrease oxygen consumption (41–43). This finding suggests that these cells might be responding to elevated reactive oxygen species (ROS) levels. The upregulation of “cellular response to stress” and “apoptotic execution phase” genes also indicates that PGRN deficient astrocytes and neurons could be under stress, that could be leading to cell death.

We found that PGRN KO astrocytes (Figure 2D), but not neurons, show upregulation of several genes involved in the glycolysis pathway (*ALDH3A2, TPI1, PKM, PGK1, ENO1, ENO2, GAPDH, PGM1, PFKP, HK1*), and the combined average expression of all genes involved in this pathway is also higher in astrocytes (Figure 2E). This suggests that astrocytes may attempt to compensate for the decreased energy production from the impaired oxidative phosphorylation by enhancing glycolysis (Warburg effect). This is also a way to support energy metabolism of neurons – lactate, produced by glycolysis in astrocytes, is taken up by neurons as an energy source (44).

Interestingly, in PGRN KO astrocytes, “stress granule assembly” was the top enriched GO biological processes term of upregulated genes (Supplementary Table S3), including *DDX6, DYNC1H1, DDX3X, YTHDF3, G3BP1, G3BP2, MAPT, BICD1*. Stress granules protects cells from oxidative stress damage, and the suppression of stress granules formation by oxidative stress promotes cell death (45). These findings suggest that PGRN KO astrocytes could regulate redox levels by inducing stress granule formation.

These findings establish a link between PGRN loss and mitochondrial dysfunction, energy metabolism, and stress response pathways, emphasizing OXPHOS as a prominent pathway affected by *GRN* deficiency.

### PGRN loss impairs mitochondrial respiration

Given the robust effect of PGRN deficiency on the expression of OXPHOS genes, we next sought to directly investigate mitochondrial function in organoids cells. We used the Seahorse XF96 Analyzer to measure mitochondrial respiration, a reliable method to assess mitochondrial function. Oligomycin (ATP-synthase inhibitor), FCCP (uncoupling agent) and rotenone/antimycin (complexes I and III inhibitors, respectively) were sequentially added to assess oxygen consumption rate (OCR) under different conditions. We observed that mitochondrial respiration, as measured by the OCR, was decreased in PGRN KO dissociated organoids (Figure 3A). Basal OCR, ATP-linked OCR, maximal respiration and reserve capacity were all significantly decreased in PGRN KO dissociated organoids (Figure 3B). The decreased reserve capacity in PGRN KO organoids indicates higher susceptibility to oxidative metabolic stress. The impaired mitochondrial respiration, with a larger proportional decrease in maximal respiration, as observed in PGRN KO organoids, can be due to several factors, such as changes in mitochondrial content or respiratory chain activity (46). Because mitochondrial mass, assayed by TOM20 expression, was not different between the two genotypes (Figure 3C, D), the observed decrease in mitochondrial respiration was most likely due to decreased activity of respiratory chain complexes I-IV. Moreover, the ATP production rate from oxidative phosphorylation was significantly decreased in PGRN KO (Figure 3E), while the ATP production rate from glycolysis (Figure 3E) and the extracellular acidification rate (ECAR) (Figure 3F), an indicator of glycolysis, did not differ between genotypes. These results indicate that PGRN loss impacts cells energy metabolism through impairment of the OXPHOS system.

**Figure 3.**
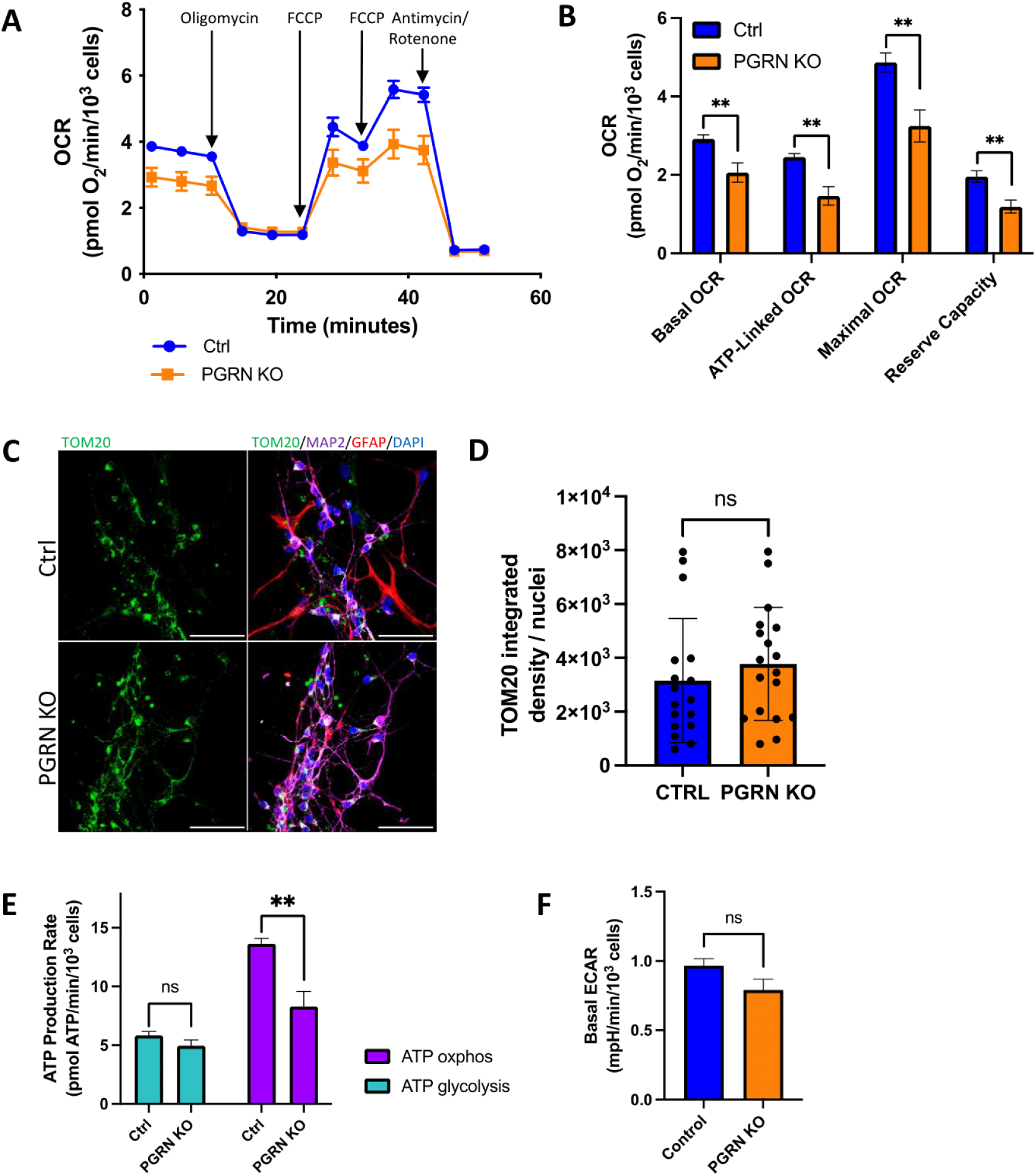
PGRN loss impairs mitochondrial respiration in human brain organoids. A – Oxygen consumption rate (OCR) was measured in PGRN KO and Ctrl dissociated organoids by the sequential addition of specific respiratory modulators. Data represent the mean ± SEM of n=6 technical replicates obtained from the dissociation of 8 organoids per genotype. Values were normalized by the number of cells counted by DAPI fluorescence. B – Respiratory parameters extracted from OCR. C – Representative images of TOM20 immunostained organoid cells from PGRN KO and Ctrl organoids, co-stained with GFAP, MAP2 and DAPI. Scale bars = 50 µm. D – Quantification of TOM20 levels in organoids, calculated as the integrated density of TOM20 fluorescence in the field of view normalized by the number of nuclei (minimum of 12 cells per field of view). E – Rates of ATP produced from oxidative phosphorylation (obtained from OCR) and glycolysis (obtained from ECAR). F – Extracellular acidification rate (ECAR) under basal conditions. Statistical significance was calculated by two-tailed unpaired *t*-test. Bars and error bars represent means ± SEM. ^∗∗^p ≤ 0.01, ns = not significant.

### PGRN loss leads to increased oxidative stress

The mitochondrial respiratory chain is the main source of intracellular reactive oxygen species (ROS), which are produced as a byproduct of electron transfer, and perturbations in the oxidative phosphorylation system can increase the production of ROS (47). Given that we observed impairment of the OXPHOS system in mitochondria, and that transcriptomics analyses suggested a dysfunctional response to oxidative stress response in PGRN KO organoids, we sought out to investigate intracellular ROS. We used CellROX green, a probe that exhibits bright fluorescence upon oxidation by ROS and localizes to the nucleus and mitochondria. CellROX labelling demonstrated a significantly higher accumulation of ROS in the PGRN KO organoid cells as opposed to controls (Figure 4A, B).

**Figure 4.**
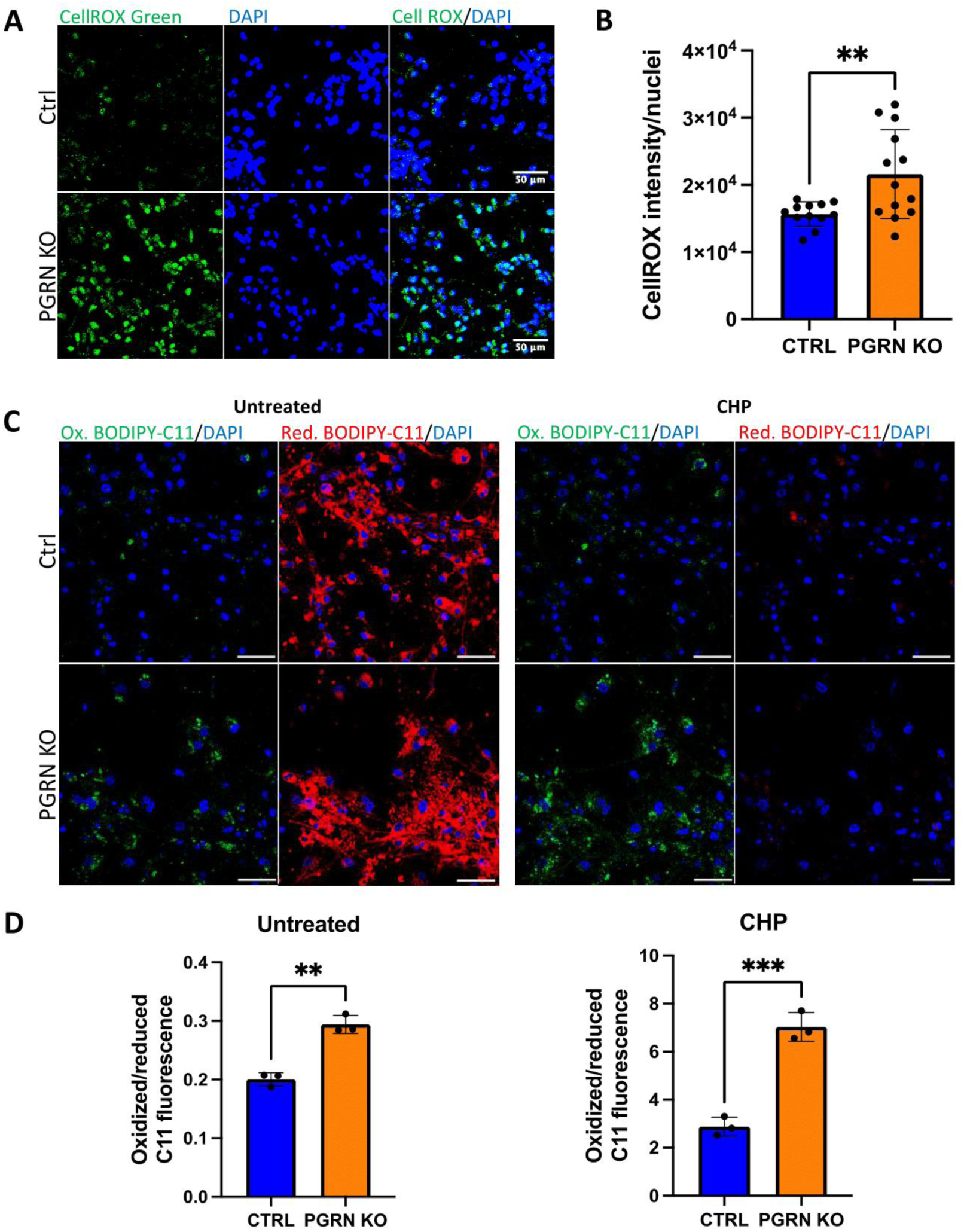
PGRN KO brain organoids show increased cellular ROS and lipid peroxidation. A – Representative images of cellular ROS in PGRN KO and Ctrl dissociated organoids by CellROX Green staining. B – Quantification of CellROX Green in organoids, calculated as the integrated density of CellROX fluorescence in the field of view normalized by the number of nuclei. C – Representative images of lipid peroxidation in untreated PGRN KO and Ctrl dissociated organoids by BODIPY 581/591 C11 staining. Ox. BODIPY-C11 (oxidized) represents the levels of lipid ROS, while Red. BODIPY-C11 represents the level of staining with the probe (reduced or unoxidized). Scale bars = 50 µm. D – Quantification of lipid peroxidation levels in untreated (left) and CHP-treated (right) dissociated organoids, calculated as the ratio of oxidized/reduced BODIPY-C11 mean fluorescence intensity in the ROI of field of view. Each point represents an average of ∼65 fields of view. Statistical significance was calculated by two-tailed unpaired *t*-test. Bars and error bars represent means ± SD. ^∗∗^p ≤ 0.01, ^∗∗∗^p ≤ 0.001.

Excessive ROS can cause oxidative damage on membrane lipids, DNA and proteins. Lipid peroxidation is the oxidative degradation of lipids, and its products (reactive aldehydes and lipid radicals) can further damage lipids and proteins in a chain reaction, resulting in the formation of lipofuscin – a lipid-protein aggregate and one of the hallmarks of NCL (48–50). Therefore, we investigated if PGRN loss alters cell lipid peroxidation levels (Figure 4C, D). We used BODIPY 581/591 C11 as a live-cell reporter for lipid peroxidation. BODIPY C11 localizes in lipophilic membrane structures, and its oxidation shifts the fluorescent emission peak from ∼590 nm (red, reduced) to ∼510 nm (green, oxidized). Under baseline conditions, PGRN KO cells showed a significant increase in lipid peroxidation (ratio of oxidized/reduced BODIPY C11 fluorescence) compared to controls (Figure 4C, D, left). After three hours of treatment with cumene hydroperoxide (CHP), a well characterized oxidant and lipid peroxidation inducer (51), lipid peroxidation levels increased considerably in both genotypes, and the increase was significantly higher in PGRN KO cells (Figure 4C, D, right). These results suggest that PGRN loss leads to increased basal oxidative stress, indicated by increased nuclear, mitochondrial and lipid-derived ROS levels, and increased cellular vulnerability to induced oxidative stress.

Interestingly, we did not find lipofuscin buildup in PGRN KO organoids (data not shown), which contradicts findings from other studies that have reported lipofuscin accumulation in *GRN*^-/-^ iPSC-derived neurons (22,23). This suggests that, in our model, lipofuscin accumulation could be a later event because of increased oxidative stress, as it’s been shown in aged *GRN*^-/-^ mouse models (52).

### PGRN loss sensitizes cells to ferroptosis

The accumulation of lipid peroxides can lead to membrane disruption and ferroptosis, a non-apoptotic type of programmed cell death characterized by iron accumulation and increased lipid peroxidation (53,54). Another hallmark of ferroptosis is impaired activity of glutathione peroxidase 4 (GPX4), an antioxidant enzyme and major inhibitor of ferroptosis that converts toxic lipid hydroperoxides to non-toxic lipid alcohols (54,55) (Figure 5A). Since our results indicate that PGRN loss leads to increased ROS and lipid peroxidation, we next investigated if PGRN deficiency is also linked to sensitivity to ferroptosis. A list of ferroptosis suppressors and driver genes was retrieved from FerrDb (56), a database of ferroptosis regulators, and used to screen for ferroptosis-related DEGs. Differential gene expression analysis showed that in PGRN KO glutamatergic neurons (CN1, CN3, CN2) several ferroptosis-suppressor genes were downregulated (*TXN, FTL, FTH1, TMSB4X, CISD1, CISD2, CHMP5, PARK7, SOX2, BEX1, GPX4*) (Figure 6B). Among these genes, only *FTH1* and *CISD2* were not downregulated in CN2. *BEX1*, that encodes brain-expressed X-linked protein 1, the most upregulated gene in this gene list, has been shown to protect cancer cells from ferroptosis (57). Thioredoxin (*TXN*) inhibits ferroptosis by regulating GPX4 and GSH (58–60), and it promotes ferroptosis in cancer cells when suppressed (58–60). *FTL* and *FTH1* encode the light and heavy chain, respectively, of ferritin, a cytosolic iron storage protein that functions as an iron buffer, and promotes increased Fe^2+^ intracellular levels and ferroptosis when downregulated (61–64). *CISD1* and *CISD2* encode for iron-sulphur cluster containing proteins that regulate mitochondrial iron and ROS metabolism, and knockdown of *CISD1* or *CISD2* induces ferroptosis and mitochondrial iron overload (65–67). In PGRN KO neurons (CN - glutamatergic neurons and IN - GABAergic interneurons), *GPX4,* a master regulator of ferroptosis, was the second most significantly downregulated gene within ferroptosis suppressor genes, and we confirmed its downregulation at the protein level in mature (MAP2+) neurons (Figure 5C, D).

**Figure 5.**
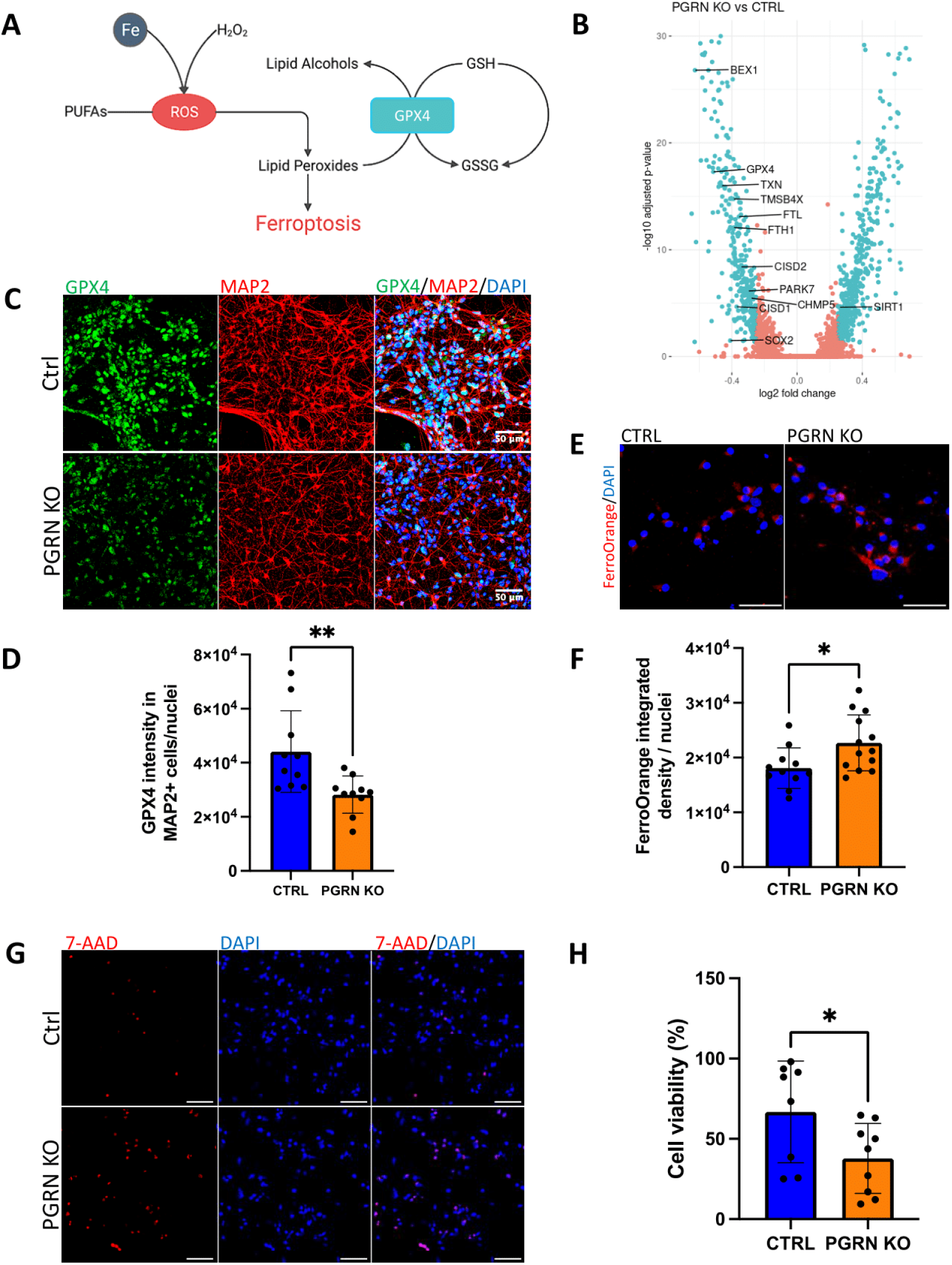
Loss of progranulin sensitizes cells to ferroptosis. A – The production of ROS can cause lipid peroxidation, which in turn induces ferroptosis. Ferrous iron (Fe^2+^) reacts with hydrogen peroxide (Fenton reaction), which generates ROS that oxidizes polyunsaturated fatty acids (PUFAs) and results in membrane damage and cell death (ferroptosis). Glutathione peroxidase 4 (GPX4) is the central enzyme regulating ferroptosis. GPX4 converts gluthatione (GSH) into oxidized glutathione (GSSG) and reduces lipid peroxides to non-toxic lipid alcohols. Diagram reprinted from *Rockland Immunochemicals,* 2023, retrieved from rockland.com/resources/the-ferroptosis-pathway. B - Volcano plot of differential gene expression results of PGRN KO versus Ctrl glutamatergic (CN1, CN2, CN3) neurons, with differentially expressed negative regulators of ferroptosis highlighted. Within the depicted genes, only *FTH1* and *CISD2* are not significantly downregulated in CN2. 925 DEGs (green) were identified using the Wilcoxon test with a log2FC cutoff of (abs)0.25 and p-adjusted value <0.05. Ferroptosis suppressor genes were collected from the ferroptosis database FerrDb v2. C – Representative images of PGRN KO and Ctrl dissociated organoids labeled with GPX4 and MAP2. D – Quantification of GPX4 in MAP2-positive cells, calculated as the integrated density of GPX4 fluorescence in MAP2+ cells in the field of view and normalized by the number of nuclei. E – Representative images of live PGRN KO and Ctrl organoids labeled with FerroOrange and DAPI. F – Quantification of FerroOrange in organoids, calculated as the integrated density of FerroOrange fluorescence in the field of view normalized by the number of nuclei. G – Representative images of PGRN KO and Ctrl organoids treated with 100 uM cumene hydroperoxide for 2 hours, and labeled with 7-AAD and DAPI. H – Cell viability based on of 7-AAD-positive ferroptotic cells after 2 and 3 hours of CHP treatment. The extent of cell death was determined based on counting 7-AAD positive dying cells in randomly chosen regions. Each field of view contains 90 +/- 31.5 cells. Statistical significance was calculated by two-tailed unpaired *t*-test. Bars and error bars represent means ± SD. ^∗^p ≤ 0.1, ^∗∗^p ≤ 0.01. Scale bars = 50 µm.

Ferrous iron (Fe^2+^) accumulation is another hallmark of ferroptosis (53). Iron can induce ferroptosis by activating lipid peroxidation enzymes or by directly generating ROS through the Fenton reaction (68,69). To investigate the overall levels of ferrous iron, we used FerroOrange, a fluorescent probe that enables live-cell fluorescent imaging of intracellular Fe^2+^. We found that PGRN KO organoids have significantly higher ferrous iron content compared to controls (Figure 5E, F). Transcriptomic analysis showed that neurons expressed lower levels of iron transport and uptake genes (Figure 2C). Looking at these results closely, we found that this difference was driven mainly by glutamatergic neurons. Given that glutamatergic neurons showed dysregulation of iron-related and ferroptosis-suppressor genes, we also investigated ferrous iron content in neurons by co-labeling live organoid cells with FerroOrange and NeuO, a fluorescent probe that selectively labels neurons. We found that NeuO-positive cells also express higher levels of ferrous iron (Supplementary Figure S3).

To determine if progranulin loss impacts ferroptotic cell death, we induced ferroptosis in organoids by exposing them to CHP. After two and three hours of treatment, cell death in dissociated organoids was assessed with 7-AAD staining. Cumene hydroperoxide induced dramatic overall cell death in both genotypes, but PGRN KO dissociated organoids showed a significant increase in relative cell death when compared to controls (Figure 5G, H).

## Discussion

Our study reveals that PGRN loss in human organoids resulted in pronounced mitochondrial dysfunction. This is evident from the downregulation of ETC genes and the subsequent reduction in oxygen consumption – an accurate measure of mitochondrial function. These data suggest impaired respiratory chain activity resulting from the lower expression of ETC genes. Mitochondrial dysfunction is a common feature in several neurodegenerative diseases, including AD, PD, ALS, Leigh syndrome, and multiple forms of NCL (70–77). These disorders often exhibit mitochondrial defects such as decreased mitochondrial membrane potential and altered respiratory chain enzyme activities. In PGRN-deficient models and cells, including human FTD-GRN neurons (78), PGRN-deficient neural progenitor and neuroblastoma cells (37,79), and mouse brains (80), mitochondrial respiration is decreased and mitochondrial genes and proteins are downregulated. More recently, it has been shown that the impact of PGRN loss on mitochondrial function in mouse models varies by genetic background, with compensatory upregulation of mitochondrial function in some strains but not others, and that compromised mitochondrial function could worsen the phenotypes associated with PGRN loss (81). Our findings indicate that PGRN loss results in excessive cellular ROS production, likely as a consequence of impaired OXPHOS and a deficient antioxidant response. PGRN has been shown to protect against oxidative stress (82,83), and excessive ROS have been reported in other progranulin-deficient disease models such as human neuroblastoma cells (37), and mouse microglial cells (84). Moreover, oxidative stress is a key contributor to a number of neurodegenerative disorders, including AD, PD, ALS and NCL (85,86). Together, our observations reinforce the notion that PGRN deficiency may converge on common pathogenic mechanisms involving mitochondrial dysfunction and redox imbalance.

A key early response to such oxidative stress is mitophagy activation; however, PGRN deficiency has been linked to impaired autophagy and mitophagy (36–40). Nevertheless, we did not detect a significant change in mitochondrial mass as assessed by TOM20 staining. This could reflect compensatory upregulation of mitochondrial biogenesis or cell type–specific differences in mitophagic activity, as previously proposed (31). Further analysis using additional mitochondrial markers and functional assays will be necessary to clarify whether the observed mitophagy impairment results in accumulation of dysfunctional mitochondria or reflects adaptive remodeling of mitochondrial turnover.

When ROS surpass the capabilities of antioxidant defense mechanisms, excessive ROS can trigger lipid peroxidation, particularly targeting polyunsaturated fatty acids in cell membranes. Consistent with this mechanism, we observed elevated lipid peroxidation in PGRN KO organoids, paralleling findings in other neurodegenerative diseases. Oxidative damage to lipids can lead to the formation of lipofuscin, a hallmark of NCL and a phenomenon also observed in PGRN haploinsufficiency (Ward et al., 2017) and other neurodegenerative disorders such as Huntington’s disease, AD and PD (50). The elevation in lipid peroxidation, together with increased ROS production and dysregulation of a number of antioxidant genes, reinforces the notion that PGRN deficiency exacerbates cellular vulnerability to oxidative stress and contributes to redox imbalance–driven neurodegeneration.

Our findings on increased susceptibility of PGRN KO cells to ferroptotic cell death add another layer of complexity to the role of PGRN in neurodegeneration. Ferroptosis is associated with a diverse array of neurodegenerative disorders, such as AD, PD and ALS (87–89). Additionally, PGRN has been shown to protect mouse brains from ferroptosis induced by cerebral ischemia (90,91) and the neurotoxin MPP^+^ (83).

We also showed that PGRN KO organoids accumulate ferrous iron, consistent with previous reports showing that knockdown of ETC genes increases labile iron (92). Mitochondria are central to iron metabolism, as they contain up to 20-50% of cellular iron, are the sole site of heme synthesis and the primary generator of iron-sulfur (Fe-S) clusters. Impaired ETC function can destabilize these co-factors and result in release of ferrous iron (92,93). The observed downregulation of GPX4 and other ferroptosis-suppressor genes in PGRN KO neurons, particularly in glutamatergic neurons, highlights the potential role of PGRN in modulating GPX4-mediated ferroptosis. Given that PGRN contains multiple cysteine granulin motif (94), its loss may disrupt cysteine and redox homeostasis. Cysteine depletion can induce ferroptosis by causing a direct inhibition of GSH and GPX4 protein expression (95), and disruption of lysosomal cystine efflux attenuates ATF4 induction and triggers ferroptosis (96).

As summarized in the graphical abstract, the interplay between mitochondrial dysfunction, oxidative stress, and ferroptosis is complex, and these mechanisms most likely collectively contribute to neurodegeneration in the context of PGRN deficiency. Additionally, PGRN plays a major role in the regulation of autophagy and lysosomal function (12,36,97). PGRN loss results in downregulation of ETC genes, either directly or as part of an adaptive response to oxidative stress, which can lead to mitochondrial dysfunction, and overproduction of reactive oxygen species (ROS) due to an incomplete reduction of oxygen in the ETC. Dysfunctional mitochondria can also disrupt lysosomal function, causing the buildup of damaged cellular components, which in turn may amplify mitochondrial stress. Defective lysosomal function leads to the accumulation of waste, inducing oxidative stress. Moreover, lipofuscin, found in dysfunctional lysosomes, sequesters iron, contributing to ferroptosis (92). Oxidative stress could further impair both mitochondrial and lysosomal functions. The accumulation of damaged cellular components can overwhelm lysosomes, and oxidatively damaged mitochondria could intensify ROS production. Oxidative stress is closely associated with ferroptosis, as it can trigger lipid peroxidation, initiating this form of cell death. Additionally, the buildup of iron catalyzes ROS formation, which can result in lipid peroxidation and ferroptosis.

Our findings provide valuable insights into the molecular mechanisms underlying PGRN deficiency. The convergence of mitochondrial dysfunction, oxidative stress and increased vulnerability to ferroptosis highlights the complex interplay of these processes in neurodegeneration caused by PGRN loss. Although we did not find differences in cell numbers between PGRN KO and control organoids, these features may represent early pathogenic events. Further investigation into the precise molecular pathways linking PGRN deficiency to these cellular abnormalities could be crucial for developing targeted therapeutic strategies for NCL and FTD caused by GRN loss-of-function mutations.

## Methods

### Human iPSC lines and culture

PGRN KO (*GRN*^-/-^) and Ctrl (*GRN*^+/+^, isogenic control) human iPSC were both generated in the WTC11 background. Ctrl contains a TdTom under *TMEM119,* and it was generated from WTC11. PGRN KO iPSC line was generated by Dr. Bruce R. Conklin (Gladstone Institutes, UCSF) (21) by mutagenizing WTC11 iPSC by dual Cas9 D10A nickase in the exon 12, which resulted in a 10 base pair deletion in one allele, and a 7 bp insertion plus a 1 bp mutation in the second allele (Figure 1A). Human iPSCs were cultured in mTeSR Plus medium (STEMCELL Technologies) in 6-well plates (Corning) coated with Matrigel (Corning). Medium was changed daily, and cells were passed with ReleSR (STEMCELL Technologies) when 70-80% confluent.

### Human forebrain organoids generation

For the generation of human forebrain organoids, we followed previously described protocols (20,31) with a few modifications. On day 0, hiPSC were dissociated into single cells with Accutase (STEMCELL Technologies), then seeded into AggreWell 800 Microwell plates (STEMCELL Technologies). 3 x 10^6^ cells were seeded per well in mTeSR Plus medium plus 10 nM ROCK inhibitor. After 24h, embryoid bodies were collected from the wells and transferred to a 10 cm dish treated with Anti-Adherence Rinsing solution (STEMCELL Technologies) in Neural Induction Medium (NIM), containing DMEM/F12 (Gibco) as the base, plus 1X GlutaMAX Supplement (Gibco), 1X MEM Non-Essential Amino Acids (Gibco), 20% KnockOut Serum Replacement (Gibco), 1x Antibiotic-Antimycotic (Gibco), 0.1 mM 2-mercaptoethanol, 5 uM dorsomorphin (Sigma), 10 uM SB-431542 (Tocris), and 2.5 uM XAV-939 (Tocris).). Human dorsal forebrain organoids (hCO) and ventral forebrain organoids (hVO) were fed daily with NIM from days 1-5, and 5 µM IWP-2 was added to medium on days 4-6 only for hVO. On day 6, organoids were transferred to Neural Differentiation Medium (NM), containing Neurobasal-A (Gibco), B-27 supplement without vitamin A (Gibco), 1x GlutaMax, and 1x Antibiotic-Antimycotic, supplemented with 20 ng/ml EGF (R&D Systems) and 20 ng/ml FGF2 (R&D Systems). hCO organoids were fed daily with NM plus FGF2 and EGF from days 6-15, and every other day from days 16-24. For hVO organoids, NM was also supplemented with 5 uM IWP-2 (Selleckchem) from days 6-11; 5 uM IWP-2, 100 nM SAG (Selleckchem) and 100 nM RA (Sigma) from days 12-14; 5 uM IWP-2, 100 nM SAG, 100 nM RA and 100 nM AlloP (Selleckchem) on day 15; and 5 uM IWP-2, 100 nM SAG, and 100 nM AlloP from days 16-24. Medium was changed daily for hVO from days 6-24. For both hCO and hVO, NM was supplemented with 20 ng/ml BDNF (Peprotech) and 20 ng/ml NT3 (Peprotech) from days 25-43, with medium changes every other day. From day 43 on, hCO and hVO organoids were maintained in NM without additional supplements and fed every 4 days. For functional experiments, only hCO were used. For scRNA-seq, hCO and hVO were fused at D60.

### Dissociation of organoids

Organoids were dissociated utilizing the Worthington Papain Dissociation System (Worthington Biochemical). Briefly, each organoid was incubated at 37°C for 30 min in a solution of papain plus DNase, after which they were triturated every 10 min with a 1000 ul pipette tip until the whole tissue was dissociated (about 4-5 cycles of trituration). The single cell solution was then mixed with an albumin-ovomucoid inhibitor solution plus DNase and pelleted with centrifugation at 300 RCF for 5 min. Cell were then used for scRNA-seq library construction or functional experiments. Organoids used for scRNA-seq were 6 months old, and for functional experiments, 5-7 months old. For functional experiments, at least 5 organoids were pooled for each dissociation. Cells were resuspended in DMEM/F12 plus 10% FBS, and plated in glass-bottom 8-well chambers (IBIDI) coated with 50 ug/ml PDL, at a density of approximately 200k cells/cm^2^. On the next day, medium was replaced with BrainPhys Neuronal Medium (STEMCELL Technologies) supplemented with 1% N2 Supplement-A, 2% NeuroCult SM1 Neuronal Supplement, 20 ng/ml GDNF, 20 ng/ml BDNF, 1 mM dibutyryl-cAMP and 200 nM ascorbic-acid (supplements were purchased from STEMCELL Technologies), plus 1x Antibiotic-Antimycotic. Cells were maintained in BrainPhys and half medium was changed every 5 days. Experiments were performed within 14 and 21 days after plating.

### Single-cell RNA Sequencing (scRNA-seq)

#### Sample processing and library preparation

For scRNA-seq, we utilized 3 fused (hCO + hVO) organoids of each genotype - PGRN KO and Ctrl, at age 6 months. Organoids were dissociated into single cells as described above. After dissociation, cells were resuspended in a PBS + 0.04% BSA solution, filtered through a 70 um Flowmi cell strainer (Sigma), and cell concentration and viability were determined with trypan blue. The average number of cells per organoid was 3.39 x 10^5^ +/- 7.99 x 10^4^ cells, and cell viability was >95% for all organoid samples. Cells were then labelled with Cell Multiplexing Oligos from the 10x Genomics 3’ CellPlex Kit (10x Genomics), which allowed cells from different samples to be pooled together prior to loading into a 10x Genomics chip. The 6 samples were pooled together, and about 43,000 cells were loaded into ChipG in order to reach a targeted cell recovery of 30,000 cells. Gel In-Bead Emulsion (GEMs), and 3’ Gene Expression and Cell Multiplexing Libraries were generated using the Chromium Next GEM Single Cell 3’ GEM, Library & Gel Bead Kit v3.1 (Dual Index) with 3’ Feature Barcode kit for Cell Multiplexing (10x Genomics). Libraries were sequenced on the Illumina NovaSeq 6000 at the UC Davis DNA Technology Core, with a depth of ∼42k reads per cell.

#### Counts matrix generation and quality control

Demultiplexed FASTQ files were aligned to the GRCh38 genome (reference refdata-gex-GRCh38-2020-A) with CellRanger v6.1.1 using the CellRanger Multi pipeline, using default parameters. Downstream analysis was performed with Seurat 4.4 in RStudio 4.3.1. The preprocessed counts matrix was transformed into a Seurat object for each sample. Only cells that expressed more than 1,300 and less than 9,000 genes, and less than 50,000 UMIs were considered for the downstream analysis. We also removed damaged cells with miQC package (98), which combines information about mitochondrial percentage and library complexity (number of genes), by filtering out cells with 75% or higher probability of being compromised. We did not check for doublets, as doublet detection tools tend to remove cells with intermediate or continuous phenotypes. After quality control analysis, we obtained a total of 9,316 cells (1,553 +/- 284 cells per sample).

#### Data processing, clustering, and cell type identification

The UMI counts were normalized and scaled using the SCTransform function in Seurat. To correct for batch effects, we performed sample integration using the 3000 most variable genes (nfeatures) identified by SCTransform. Cells were clustered based on the first 100 principal components (PCs), chosen based on the elbow plot, and using a resolution of 0.6. Data were visualized in 2 dimensions with Uniform Manifold Approximation and Projection (UMAP) using the same PCs. Clustering using these parameters resulted in the identification of 18 clusters. Two small clusters that contained less than 10 cells in two or more organoids were removed from analyses. Cluster markers were identified using the FindAllMarkers function using the Wilcoxon Rank Sum test, log(fold change) threshold of 0.25, min.pct of 0.25, and min.diff.pct of 0.2. FDR-corrected p-values of ≤ 0.05 were considered significant. Cell-type annotation was performed manually based on established marker genes, on the PanglaoDB scRNA-seq database (99), on the gene list enrichment analysis tool Enrichr (100), and based on developing brain and organoid-specific scRNA-seq datasets (19,32,33,101).

#### Differential gene expression analysis and gene ontology (GO)/pathway enrichment analysis

Following cell type identification, each set cell cluster was subset, and differential gene expression was performed. Differentially expressed genes (DEG) for each cell cluster between PGRN KO and Ctrl organoids were identified with the FindMarkers function, using Wilcoxon Rank Sum test, 0.25 as the log2(fold change) threshold, and FDR-corrected p-value of ≤ 0.05. Ribosomal genes (RPS- and RPL-) were removed from analysis. Enrichment analysis for gene ontology (GO) terms and pathways among DEGs was performed using EnrichR (100) and Metascape (102). Gene set expression scores were calculated using the AddModuleScore function in Seurat, with default parameters.

### RT-qPCR

To quantify *GRN* mRNA, total RNA was extracted from organoids using PureLink RNA Mini Kit (ThermoFisher), and cDNA synthesized with the ProtoScript II First Strand cDNA Synthesis Kit (NEB). qPCR was performed on an Applied Biosystems QuantStudio 6 Pro Real-Time PCR System, using TaqMan probes (ThermoFisher) specific for *GRN* (Hs00963707_g1) and for *PPIA* (endogenous reference, Hs99999904_m1). Expression fold change was calculated using the ΔΔCt method.

### Cellular ROS

For reactive oxygen species (ROS) detection, we used CellROX Green (Invitrogen), a fluorescent probe used to measure mitochondrial and nuclear ROS. Dissociated organoid cells were incubated in fresh medium plus 5 uM CellROX Green reagent for 30 min at 37 °C. Cells were washed twice with PBS, fresh culture media plus 0.5 ug/ml Hoechst 33342 was added, and live cells were imaged immediately.

### Lipid peroxidation

The lipid peroxidation sensor BODIPY 581/591 C11 (Invitrogen) was used to detect *in-situ* lipid peroxidation in live cells. Dissociated organoid cells were incubated in fresh medium containing 5 μM of BODIPY C11 and 0.5 ug/ml Hoechst 33342 at 37 °C for 30 min. Cells were washed twice with PBS, fresh culture media was added, and cells were imaged immediately. Media was then replaced with fresh media plus 100 uM cumene hydroperoxide (Sigma). Cells were incubated for 3 h, then imaged again in the fileds of view. In the analysis, the oxidized/reduced fluorescence ratio of BODIPY-C11 was used to normalize for cell incorporation of the probe into membrane.

### Intracellular labeling of ferrous iron

FerroOrange was used to detect intracellular Fe^2^ in live cells. Dissociated organoids were incubated in fresh medium containing 1 μM of FerroOrange (Dojindo) and 0.5 ug/ml Hoechst 33342 at 37 °C for 30 min, and cells were imaged immediately. For assessment of FerroOrange in neurons, dissociated organoid cells were incubated with 5 uM AraC from D1 post-dissociation until the day of the assay (21 days), to eliminate mitotic cells and keep only post-mitotic neurons. Neurons were imaged with FerroOrange, 0.1 μM NeuO (STEMCELL Technologies) and Hoechst 33342.

### Cell death analysis

Analysis of dell death by ferroptosis was performed by incubating dissociated organoid cell cultures with 100 uM cumene hydroperoxide for 2-3 hours at 37°C. Cells were washed twice with PBS, then incubated with fresh media plus 10 ug/ml 7-AAD (Sigma) for 15 min at 37°C. Cells were fixed in 4% PFS for 10 min at room temperature (RT), washed 3x with PBS, then mounted with Prolong Diamond Antifade Mountant (Invitrogen).

### Immunofluorescence and immunohistochemistry of organoid cryosections and 2D cultures

Whole organoids were fixed in PFA 4% for 1h at RT, washed 3x with PBS, and then cryopreserved by overnight incubation at 4°C in a solution of 30% sucrose in PBS. Cryopreserved organoids where then embedded in Neg-50 Frozen Section Medium (Epredia), flash frozen in dry ice, and stored at -80° C. Frozen organoids were sectioned in 20µm slices in a Leica Cryostat CM1850. 2D dissociated organoid cultures were fixed with 4% PFA for 10 min at RT, then washed 3x with PBS.

For immunofluorescence, fixed organoid cryoslices and 2D dissociated organoid cultures were incubated in blocking buffer containing PBTA (0.5% BSA and 0.1% Triton X-100 in PBS) plus 5% normal goat or donkey serum for 1h at RT. Samples were then incubated with primary antibodies in blocking buffer overnight at 4°C, washed three times with PBTA, then incubated with secondary antibodies in blocking buffer for 1h at RT. Samples were washed three times with PBTA, then mounted with Prolong Diamond Antifade Mountant (Invitrogen) with or without DAPI for nuclei staining.

For immunohistochemistry, after inactivation of endogenous peroxidases (3% hydrogen in PBS), antibody specific antigen retrieval was performed, and cryosections were blocked and afterwards incubated with the primary antibody for 1 h. Sections were then incubated with rabbit anti-Goat IgG (H+L) secondary antibody, HRP (Invitrogen 31402, 1:500). Next, for detection of specific binding, the mouse and rabbit specific HRP/DAB IHC detection Kit micro-polymer (Abcam ab236466) was used which contains secondary antibodies and DAB to produce a dark brown color. Sections were then counter-stained with hematoxylin, dehydrated and cover slipped with permanent mounting agent.

The following primary antibodies were used: goat anti-PGRN (R&D Systems AF2420, 1:700), rabbit anti-SATB2 (Abcam ab34735, 1:100), rat anti-CTIP2 (Abcam ab18465, 1:200), guinea-pig anti-MAP2 (Synaptic Systems 188004, 1:500), mouse anti-TUJ1 (Sigma T8578, 1:500), chicken anti-GFAP (Abcam ab4674, 1:500), rabbit anti-TBR1 (Abcam 31940, 1:500), mouse anti-CUX1 (Abcam 54583, 1:500), rabbit anti-SOX2 (Sigma ab5603, 1:500), mouse anti-TOM20 (Santa Cruz sc17764, 1:500), rabbit anti-GPX4 (Abcam ab125066, 1:500), mouse anti-NeuN (1:1,000). The following secondary antibodies were used: goat anti-rabbit Alexa Fluor 488 (ThermoFisher A-11008), goat anti-guinea pig Alexa Fluor 594 (ThermoFisher A-11076), goat anti-rat Alexa Fluor 647 (ThermoFisher A-21247), goat anti-mouse AlexaFluor 555 (ThermoFisher A-21422), goat anti-chicken Alexa Fluor 647 (ThermoFisher A-21449), goat anti-mouse Alexa Fluor 647 (ThermoFisher A-21235). Secondary antibodies were used at a dilution of 1:1,000.

### Image acquisition and analysis

Samples were imaged using a Leica SP8 confocal microscope, and image analysis was done in ImageJ2 (Fiji) software v2.14.0. Background was subtracted from each image. TOM20, CellROX and FerroOrange fluorescence intensities were quantified by thresholding the targeted channel, and measuring the integrated density (IntDen) for the region of interest (ROI) in the field of view (FOV). IntDen was then normalized by the number of cells, equivalent to the number of DAPI-positive nuclei, in each FOV. For FerroOrange intensity analysis in neurons, the NeuO channel was thresholded and applied as a mask in the FerroOrange channel, and the mean fluorescence intensity of FerroOrange was measured in the ROI. For GPX4 intensity analysis, the MAP2 channel was thresholded and applied as a mask in the GPX4 channel. The IntDen was measured in the GPX4 channel for the ROI, and the value was normalized by the number of cells in each FOV. For quantification of BODIPY-C11, the images of oxidized and reduced BODIPY-C11 were first thresholded, and IntDen was measured for the ROIs. The ratio of oxidized/reduced BODIPY-C11 was calculated by calculating the ratio of oxidized/reduced BODIPY-C11 IntDen, in order to normalize for cell incorporation of the probe into membrane. For cell death analysis, the number of DAPI+ and 7-AAD positive were calculated for each FOV, and cell viability calculated as 1-(number of 7-AAD+ nuclei/number of DAPI+ nuclei) x 100.

### Oxygen consumption rate

The oxygen consumption rate (OCR) and the extracellular acidification rate (ECAR) were measured in a Seahorse XF96 Extracellular Flux Analyzer (Agilent Technologies). Organoids were dissociated, and a total of 3×10^5^ cells/well were seeded on a Seahorse XF96 plate and cultured for 2 weeks. On the day of the assay, cells were washed with assay medium (DMEM assay medium supplemented with 5 mM HEPES, 5 mM glucose, 2 mM glutamine, and 1 mM sodium pyruvate; pH 7.4). Compounds were injected sequentially during the assay resulting in final concentrations of 2 μM oligomycin (Port A), 0.75 μM FCCP (Port B), 1.35 μM FCCP (Port C), and 1 μM rotenone and 2 μM antimycin (Port D). OCR and ECAR were measured in parallel. Seahorse data were normalized to cell number by counting the number of nuclei stained with 1µg/mL Hoechst. Nuclei counts and analysis were completed on an Operetta High-Content Imaging System (PerkinElmer). ATP production rates were calculated as described by Divakaruni et al. (JBC, 2017).

## Acknowledgments

This research was funded in part by Conselho Nacional de Desenvolvimento Científico e Tecnológico (CNPq - 202991/2015-6) to C.M.C. We thank Li Gan (Cornell University), Claire Clelland and Bruce Conklin (University of California San Francisco) for kindly providing the hiPSCs used to produce the brain organoids in this study. Seahorse Extracellular Flux analysis was performed at the UCLA Mitochondria and Metabolism Core by Linsey Stiles. We acknowledge the use of the Leica confocal microscope in the NRI-MCDB Microscopy Facility at University of California, Santa Barbara, supported by NSF MRI grant DBI-1625770. We acknowledge the use of the DNA Technologies and Expression Analysis Core at the UC Davis Genome Center, supported by NIH Shared Instrumentation Grant 1S10OD010786-01.

