## Supplementary Figures for "Progranulin loss induces mitochondrial dysfunction and ferroptosis in human cerebral organoids"

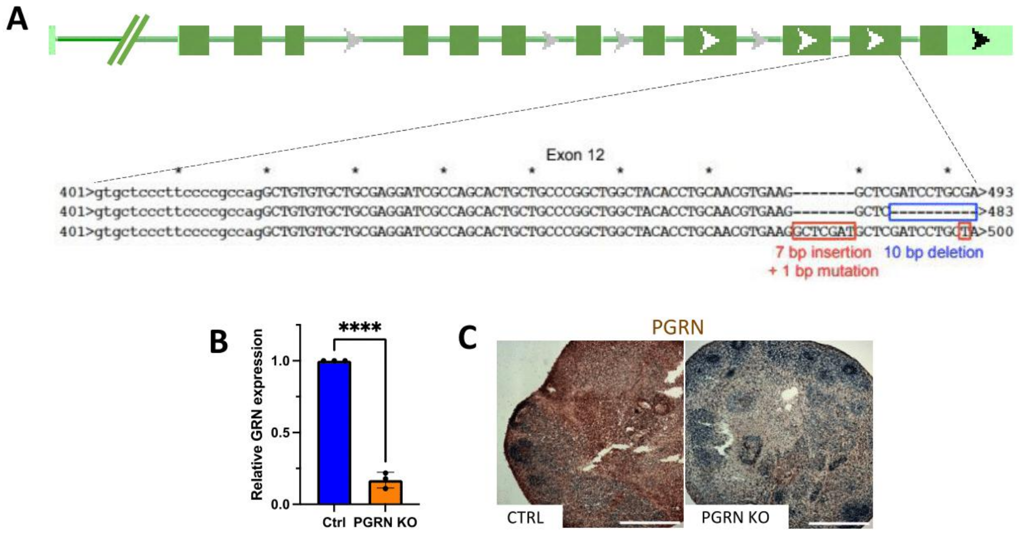


**Supplementary Figure S1 - Characterization of PGRN KO and Ctrl human brain organoids**
A – Diagram of the *GRN* gene, which contains 13 exons. The *GRN* region that was modified is depicted. The modification was generated by dual Cas9 D10A nickase in the exon 12, and resulted in a 10 base pair deletion in one allele, and a 7 bp insertion plus a 1 bp mutation in the second allele.
B – *GRN* mRNA expression in PGRN KO and Ctrl organoids. qPCR was performed using mRNA from 3 organoids from 3 different batches, for each genotype, and *GRN* expression was normalized with *PPIA*. Statistical significance was calculated by two-tailed unpaired *t*-test. Bars and error bars represent means ± SEM. *p ≤ 0.1, **p ≤ 0.01, ***p ≤ 0.001, ****p ≤ 0.0001.
C – Immunohistochemistry representative images showing expression levels of PGRN in CTRL and PGRN KO brain organoids (scale bars = 500 µm).

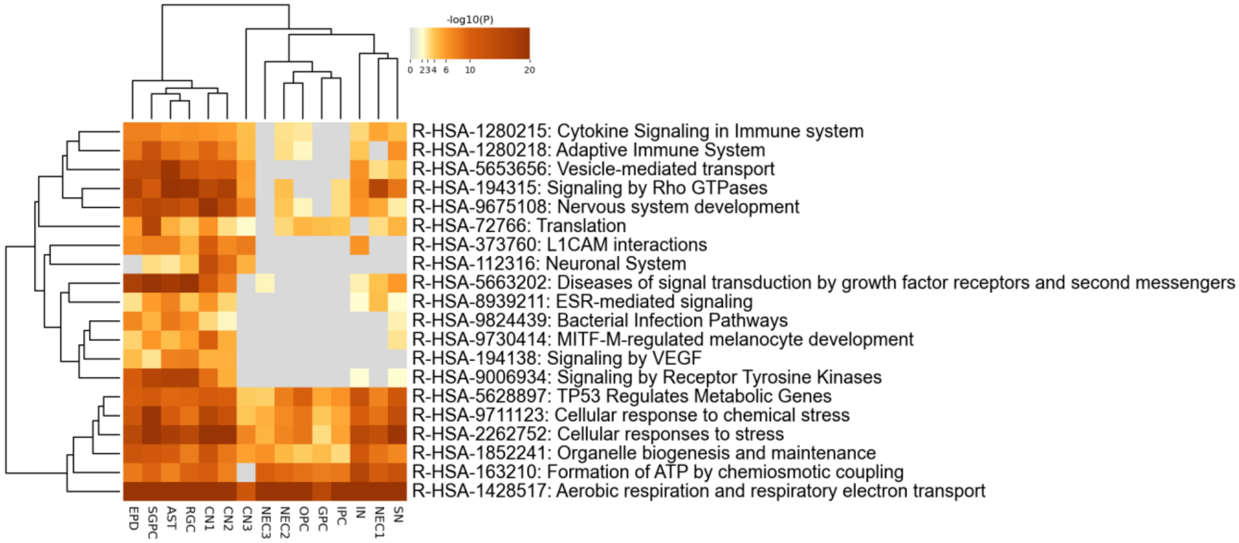


**Supplementary Figure S2. Heatmap of enriched Reactome pathways across gene-expression signatures from all cell types, colored by –log10(p-value).**

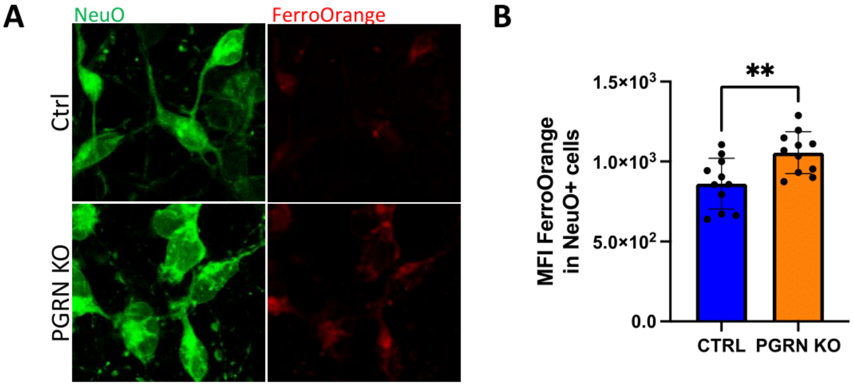


**Supplementary Figure S3 – PGRN KO neurons accumulate iron**

A – Representative images of live PGRN KO and Ctrl organoids labeled with FerroOrange and NeuO.

B - Mean fluorescence intensity (MFI) of FerroOrange in NeuO-positive cells in PGRN KO and Ctrl organoids.

Statistical significance was calculated by two-tailed unpaired *t*-test. Bars and error bars represent means ± SD. $^{**}p \leq 0.01$.
